# Does visitation dictate animal welfare in captivity? – A case study of tigers and leopards from National Zoological Park, New Delhi

**DOI:** 10.1101/2020.07.17.208322

**Authors:** Gupta Avni, Vashisth Saurabh, Sharma Mahima, Singh Randeep, Hore Upamanyu, Lee Hang, Pandey Puneet

**Affiliations:** Amity Institute of Forestry and Wildlife, Amity University, Noida, Uttar Pradesh – 201301; National Zoological Park, Mathura Road, New Delhi – 110003; Conservation Genome Resource Bank for Korean Wildlife and Research Institute for Veterinary Science, Seoul National University College of Veterinary Medicine, 1 Gwanak-gu, Gwanak-ro, Seoul – 08826, South Korea

**Author notes:** Both authors have contributed equally to this work (Gupta Avni, Vashisth Saurabh).

**Keywords:** ethogram, stereotypy, visitor effect, stress, felids

## Abstract

Zoological Parks serve a salient purpose of entertaining many visitors by housing various exclusive animal species. Big cats like tigers and leopard are among the most visited species in zoos globally. We investigated the behavioral response of the zoo-housed big cats to visitor densities and noise. We also aimed to understand the relationship between stereotypy, animal history, feeding schedules, and enclose design. The behavior of eight big cats housed in the National Zoological Park, New Delhi, was monitored using the focal sampling technique during the May and June 2019 to construct the ethograms. We also recorded the visitor density, ambient noise, for the same duration. Both species were found devoting a significant amount (>50%) of time in displaying inactive behaviors. Tigers and leopards performed stereotypic behaviors for 22% and 28% of their time, respectively. Pearson chi-square analysis revealed a significant variation of stereotypy in association with biological (age, sex, and rearing history) and captive (enclosure design) variables. Big cats’ stereotypic behaviors were found significantly influenced by the high visitor density. However, ambient noise did not impact the stereotypy of both the felid species. Visitors form an integral part of zoos, and their detrimental impact diminishes the well-being of captive animals. This study revealed that tigers and leopards in NZP display a high proportion of inactive and stereotypic behaviors. Thus, we suggest zoo authorities adopt more enclosure enrichment initiatives.

## 1. Introduction

Zoos are a means to ensure the physical and mental well-being of captive animal populations that support the existing population in the wild. Optimal animal welfare fosters comfortability, safety, ability to express innate and natural behaviors, and prevents distress. However, any environmental stressor may lead to distortion in coping with the surroundings (Morgan & Tromborg, 2007). Reduced life expectancy, diminished growth, impaired reproduction, diseases, behavior anomalies, and body damage imply sub-optimal welfare (D. M. Broom, 1991). Captive conditions are vastly different from the wild environments in terms of spatial confinement, restrictions, simplicity, control, and predictability (Morgan & Tromborg, 2007). Captivity curbs animals to elicit the complex behavior repertoire that they have evolved over the years. Zoo-housed animals necessitate mechanisms to cope up with such monotonous surroundings (Mason, 1991). The display of various forms of stereotypic behaviors suggests stress and frustration in captive conditions. Caged animals often perform repetitive, abnormal behaviors such as pacing, coprophagy, overgrooming, and head-weaving (Lyons, Young, & Deag, 1997). Such behaviors are a method to pass the time and substitute free-ranging behavior as they have no apparent function or goal (Carlstead, 1998; Hediger, 1950; Odberg, 1978). Stereotypies are supposed to deviate an individual from a typical behavior repertoire by their evident lack of purpose and have claimed to represent efforts to subsist one with unpleasant environmental conditions (Dantzer, 1991; Swaisgood & Shepherdson, 2005). These atypical behaviors indicate the suboptimal level of an animal’s psychological welfare (Boorer, 1972; Mason, 1991). Stereotypies may arise from a primary behavior pattern that caged species have eventually become motivated to perform (Holzapfel, 1939; Mason, Clubb, Latham, & Vickery, 2007).

Humans tend to leave an impact on their surrounding environment. Visitors are crucial to govern the animal welfare in captivity as they often induce alterations to behavior repertoire in captive species (Davey, 2007; Hosey, 2000). Public visiting zoological parks form relationships with captive species (Cole & Fraser, 2018). The “visitor effect” could be positive, neutral, or negative (Hosey, 2008; Hosey & Melfi, 2015). When human interactions benefit the caged animal and increase the animal’s species-specific behavior, they foster positive and healthy relationships with animals (Baker, 2004; Claxton, 2011). This results in a significant reduction in the time spent performing stereotypies and inducing natural or wild-type behaviors. There may also be certain conditions when animals become habituated to visitors due to consistent exposure and thereby exhibit no behavioral changes. On the other hand, the visitor’s unfitting behaviors can result in an adverse effect of visitation. Activities such as shouting, teasing, throwing stones, hitting, and moving in unpredictable ways can impel fear and stress response in captive animals (Cole & Fraser, 2018; Venugopal & Sha, 1993). The mere presence of human visitors yields a significant impact on the behavior of various mammalian species in zoos (Hosey, 2000). Different visitor attributes, such as presence, density, activity, noise, and proximity, can influence captive individual’s behavior and physiology (Brouček, 2014; Davey, 2005, 2007). Prevalence of stereotypy may intensify on the days of a large and noisy human audience (Dybowska, Gorecka, Grzegrzółka, Wieczorek, & Zlamal, 2008; Mallapur & Chellam, 2002; Vidal et al., 2016).

The activity pattern of an animal is an expression in response to the resources available in surroundings, and hence in zoos, animals display distinct behaviors as compared to those in the wild (Young, 2003). Behavior studies, in relation to the knowledge of species-specific behaviors in the wild, help to assess the welfare of zoo-housed animals (Keeling & Jensen, 2002). The behavior of animals reflects its first attempt to cope with sub-optimal environmental conditions and hence acts as an effective useful welfare indicator (Bashaw, Bloomsmith, Marr, & Maple, 2003; Dawkins, 1998). Studying the extent to which visitation and other captive factors influence captive specie’s behavior is a non-invasive measure. It is pivotal for suggesting better management practices for upkeep and welfare.

National Zoological Park (NZP) is one of the prominent Indian zoos. It entertains a large number of visitors each year. During 2018-19, more than 2.7 million people paid a visit to the zoo (Annual Report 2018-19). Big cats like tigers *(Panthera tigris)* and leopards *(Panthera pardus)* form a center of attraction for the most public. Visitors are more attentive and spend a long time viewing the animal when they are active and display species-specific behaviors (Altman, 1998; Bitgood, Patterson, & Benefield, 1988; Fernandez, Tamborski, Pickens, & Timberlake, 2009; Margulis, Hoyos, & Anderson, 2003). The study intended to understand the extent of stereotypic behaviors in the activity pattern of captive tigers and leopards housed at National Zoological Park, New Delhi. Additionally, it aimed to understand the relationship between visitation, ambient noise, and stereotypy.

## 2. Methodology

### 2.1. Study area and subjects

National Zoological Park, situated in India’s capital, received the status with the idea of it being the model zoo for the entire country (Agnihotri, 2012). The large-category zoo spreads over 188.62 acres of land and houses about 100 species in a total of 72 enclosures. It boasts a distinguished history of successful breeding of various animals including, tiger, brow-antlered deer, Indian rhinoceros, and Asiatic lion (Agnihotri, 2012).

We studied a total of eight individually-housed subjects (Table 1). These included four individuals each of tiger (*Panthera tigris tigris*) and leopard (*Panthera pardus fusca)*. Three of the tigers were white and one a normal variant. All three white tigers were housed in one enclosure with an arena area of 1445 sqm. The studied normal tiger was housed in an enclosure of about 858 sqm arena area. The leopards were housed in two adjacent enclosures with an arena area of about 158 sqm and 136 sqm. A pair of male and female was let out into adjacent on-exhibit enclosures together. The enclosures were enriched with logs, trees, vegetation, and water supply. Subjects studied were let out in the on-exhibit enclosure from 09:30 to 16:30 hours. The animals were fed with buffalo meat once a day in their night cells, except Fridays. Since no animals are released in the enclosure on Friday, the subjects were studied for six days a week.

**Table 1.** Subjects studied and their history (ZIMS)

| Individual | Coat Colour | Sex | Age<sup>†</sup> | Origin | Rearing History |
| --- | --- | --- | --- | --- | --- |
| Tiger (*Panthera tigris*) |  |  |  |  |  |
| Vijay | Mutation | Male | 12 | Captive (Delhi zoo) | Parent |
| Kalpana | Mutation | Female | 12.4 | Captive (Delhi zoo) | Parent |
| Karan | Normal | Male | 6 | Captive (Mysore zoo) | Parent |
| Sita | Mutation | Female | 4.3 | Captive (Delhi zoo) | Unbred |
| Leopard (*Panthera pardus*) |  |  |  |  |  |
| Tejas | Normal | Male | 6.8 | Wild (Uttarakhand) | - |
| Babli | Normal | Female | 2.25 | Wild (Jammu) | - |
| Bunty | Normal | Male | 7.9 | Wild (Chhattisgarh) | - |
| Bunty | Normal | Female | 8.8 | Wild (Chhattisgarh) | - |

<sup>†</sup> = as on 30.07.2019

### 2.2. Activity Budgeting

The study was conducted for 208 hours, during the summer months of May and June in 2019. A pilot study of three days helped to enlist all the behaviors performed by the two species (Supplementary Table 1). Each individual was observed for 6 hours 30 minutes per day, for 4consecutive days. On Fridays, the animals are fasted and kept in off-exhibit cells; hence they were not studied. A focal animal behavioral sampling at 1-minute intervals was used to construct an ethogram of the big cats (Altmann, 1974). The activities performed were classified into three categories - a) active, b) inactive, c) stereotypic, for comparison and analysis (Table 2). Active behaviors include activities like walking, eating, drinking, chewing, and playing. Activities like sleeping, sitting, and resting are classified under inactive behaviors. Stereotypic behaviors were performed in various forms, including pacing, skip-pacing, and tail or toe sucking. The behavior was considered pacing when animal covered three or more traverses of a definite path (Forthman & Bakeman, 1992). The enclosures were divided into different zones – middle, edges, and visitor areas. Zone utilized for each state of behavior by animals was also recorded.

**Table 2.**
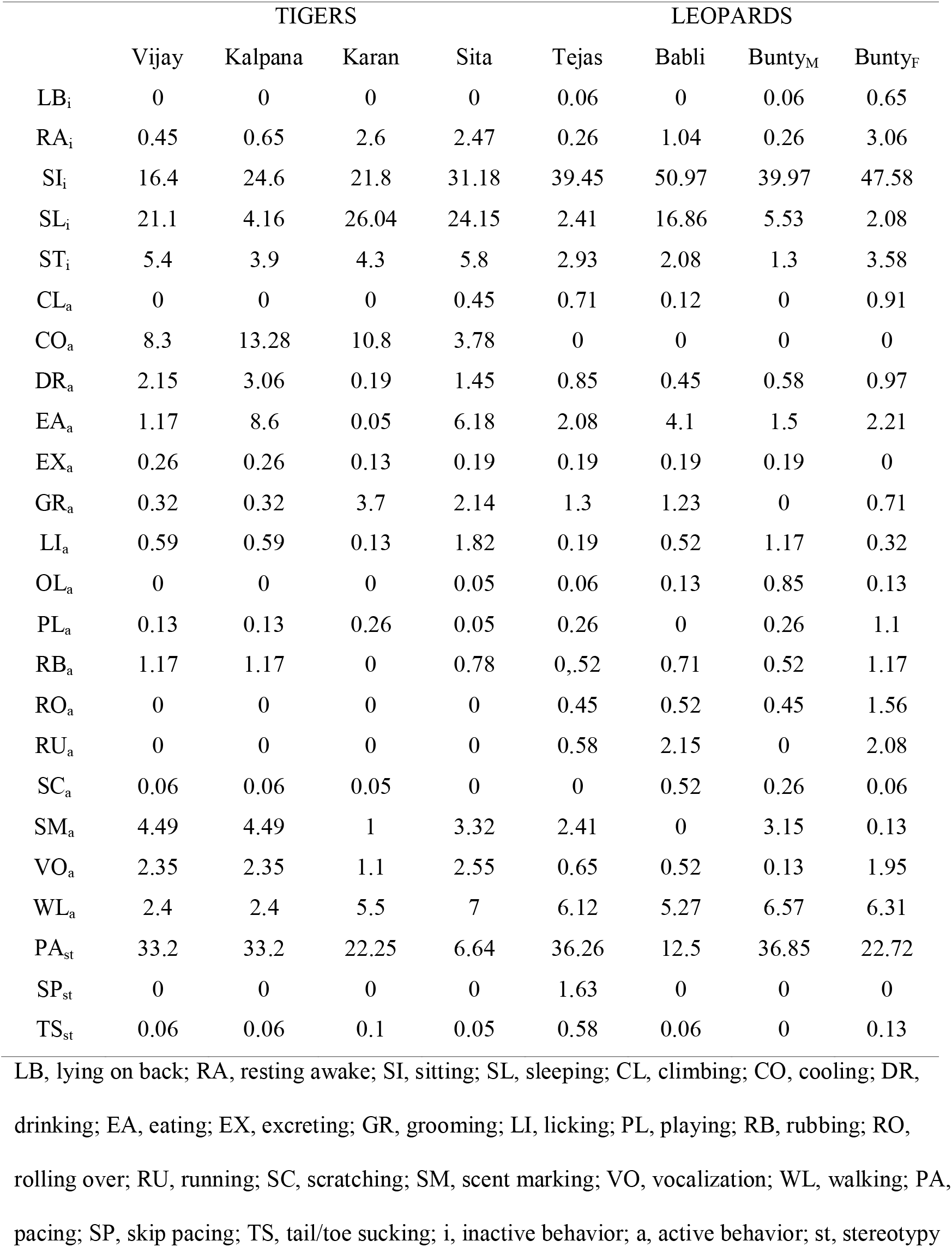
Percent of time spent in displaying different behaviors by tigers & leopards at NZP

### 2.3. Biological and Captive Factors

The history of all the studied individuals was obtained from Zoo Aquarium Animal Management Software (ZIMS). Aspects such as age, sex, birthplace, origin, time spent in captivity, and rearing history were recorded (Table 1). Age was classified as young/adult; sex as male/female; origin as wild-born/captive-born; and rearing history as bred/unbred. Association of these variables was evaluated with stereotypic levels of all tigers and leopards.

### 2.4. Visitation

Visitation and behavior data were collected simultaneously yet independently. Visitor aspects considered were – visitor density (crowd size) and ambient noise. Visitor density was calculated by counting the number of visitors standing at the visitor area around the enclosures. Ambient noise was measured in Decibels (dB), using digital noise meter (970P Meco Digital Sound Level Meter).

### 2.5. Statistical Analysis

Ethogram was constructed by calculating the frequency of each behavioral activity, converted to the proportion (or percentage) of time devoted to the particular behavior. Time spent in performing stereotypic and non-stereotypic behaviors was calculated. These were represented as strings of 0’s and 1’s, where 1 denoted presence of stereotypy and 0 denoted absence of stereotypy. Data were recorded and analyzed in Microsoft Excel 2019. The average visitor density per minute and average ambient noise per minute at the cage was calculated. The readings were found to deviate from a normal curve; hence, non-parametric tests were performed. Statistical analysis was made using the software IBM SPSS for Windows (Version 23). In all the tests, *p*-value (α) was defined at the value of 0.05 to establish statistical significance. Pearson’s chi-square association test was performed to understand the relationship between animal history and stereotypic behaviors. Binary Logistic Regression model was performed to understand the effect of visitation (visitor density and ambient noise) on stereotypy. Visitation characteristics were considered as independent variables, while animal behavior was taken as the dependent variable.

## 3. Results

### 3.1. Activity budgeting

The activity budget of tigers (n=4) revealed that they devoted a significant amount of time displaying inactivity (49±13%), of which, sitting was the most common (23±6%) (Figure 1 and Table 2). Animals utilized the middle area and edges of enclosures for inactive behaviors. They devoted 29±8% of their time to active behaviors (Figure 1). Cooling was the most performed behavior (9±4%), followed by walking (5±2%). The middle, enriched zones of the enclosures were generally utilized for active behaviors. Tigers performed stereotypy for 22±11% of their time. Pacing (22±11%) and tail or toe sucking (0.1%), were the two forms of stereotypic behaviors. The pacing was predominantly performed toward the edges of the enclosure.

**Figure 1.**
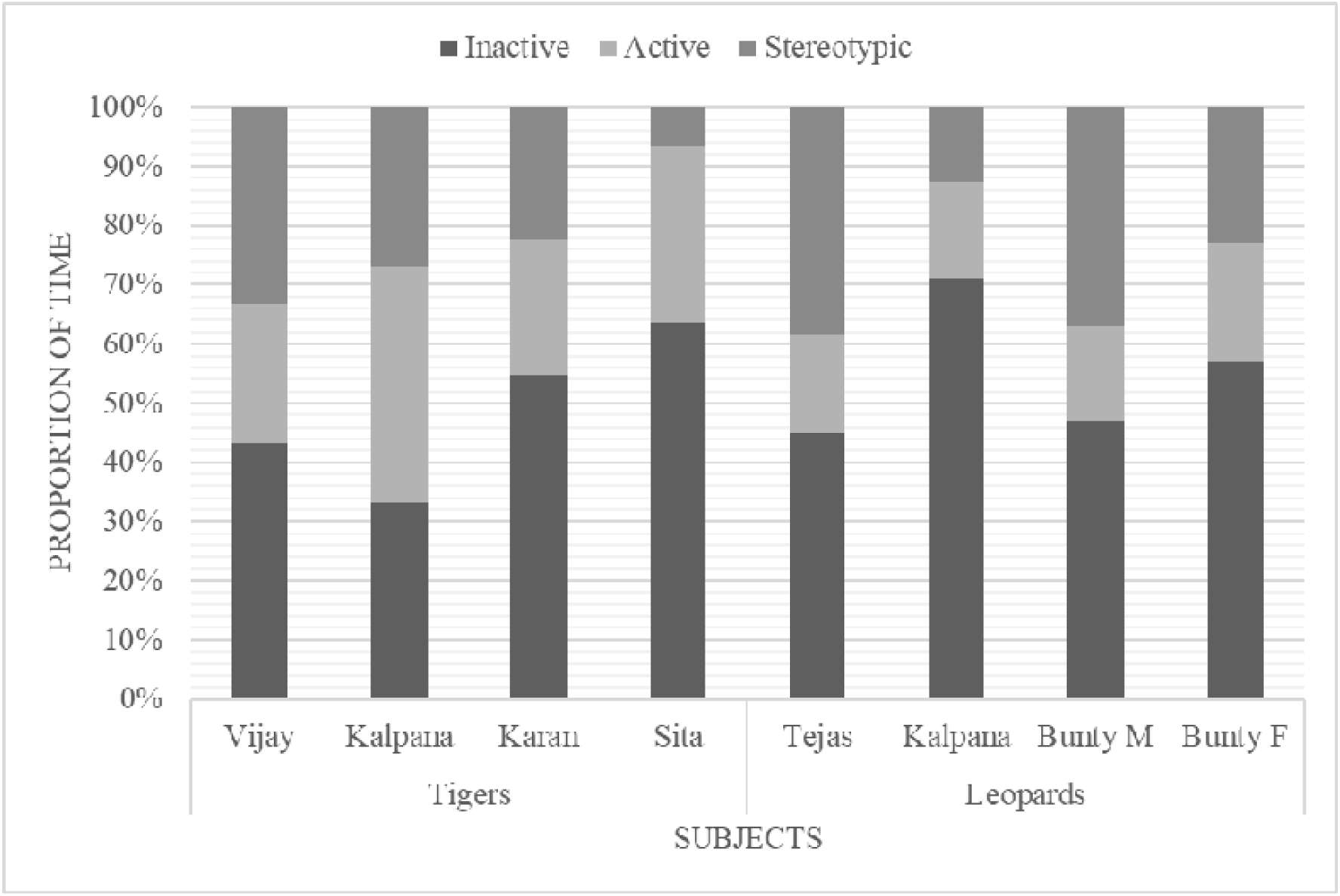
Proportion of time spent by each subject of tigers and leopards in performing behaviors – a) inactive, b) active, c) stereotypic

Like tigers, leopard subjects also devoted a considerable amount of time to inactive behaviors (55±12%, Figure 1). Sitting was the most common inactive behavior (41±5%, Table 2). The middle and rear areas of the enclosure were generally utilized for inactivity. They display active behaviors only for about 17±2% of their time in on-exhibit enclosure. The highest amount of time was devoted to walking (6±1%) of active behaviors (Table 2). Enriched zones of the enclosures were found to be utilized during the active periods. Leopards spent about 28±13% of their time in displaying stereotypy. Unlike tigers, leopards display a more varied form of stereotypy, including pacing (27±12%), skip-pacing (0.4%), and tail or toe-sucking (0.2%) (Table 2). They paced toward the edges and visitor zone of the enclosure.

### 3.2. Biological and Captive Factors

Chi-square test analysis indicated the difference in stereotypy among the two sexes (Table 3). Males exhibited high level of stereotypy (tiger=28±8%, leopard=38±1%) as compared to females (tiger=17±14%, leopard=18±7%). Analysis of stereotypy with respect to rearing history revealed a significant difference (□^2^=39.51; *p*<0.05) in tigers as parents (n=3) displayed more stereotypic behaviors (28±5%) than unbred animals (7%). In both species, adults performed much more stereotypy in comparison to young (Table 3). However, the relationship between age and stereotypy was found significant only in tigers (□^2^=30.14; *p*<0.05). Tigers housed in different enclosures did not reveal significant variation in stereotypic levels (□^2^=0.07; *p*>0.05). Leopards housed in an enclosure with more viewing area (n=2) performed high levels of stereotypy (31±11%) in comparison to those housed in an enclosure with limited viewing area (25±17%). The relationship was statistically significant (□^2^=36.58; *p*<0.05), suggesting the influence of enclosure type on the behavior repertoire.

**Table 3.** Pearson’s Chi-Square analysis to identify association of stereotypic behaviors displayed by captive tigers and leopards with biological variables (sex, rearing history, and age) and captive variable (enclosure design) at NZP

|  | Pearson Chi-Square of<br>stereotypy w.r.t. | Value | df | Asymptotic<br>Significance | Exact<br>Significance |
| --- | --- | --- | --- | --- | --- |
| Tigers | Gender | 13.17 | 1 | 0.00 | 0.00 |
|  | Rearing history | 39.51 | 1 | 0.00 | 0.00 |
|  | Age | 30.14 | 1 | 0.00 | 0.00 |
|  | Enclosure | 0.07 | 1 | 0.79 | 0.83 |
| Leopards | Gender | 36.58 | 1 | 0.00 | 0.00 |
|  | Age | 0.601 | 1 | 0.21 | 0.23 |
|  | Enclosure | 36.58 | 1 | 0.00 | 0.00 |

### 3.3. Visitation Effect

The average density of the audience was found higher around tiger enclosures (31 humans/minute) compared to leopards (13 humans/minute). However, the average noise level remains almost the same around the enclosures of both species (tiger=64 dB and leopard=62 dB). Binary logistic regression model was able to distinguish between the effect of stereotypy and visitation on both species □^2^ =4.17 for tigers, □^2^ =8.14 for leopards; df=1; *p*<0.05) (Table 4). The level of stereotypy gets increased when they are encountered with large crowds of audiences. However, the tiger and leopard behavior repertoire remained unaffected by the ambient noise as the relationship was non-significant (□^2^=4.17 for tigers; □^2^=8.14 for leopards; df=1; *p*>0.05) (Table 4).

**Table 4.** Binary logistic regression to understand visitation effect on tigers and leopards at NZP

|  |  | B | S.E. | Wald | df | Significance | Exp<br>(B) | 95% C.I. for<br>Exp (B) |  |
| --- | --- | --- | --- | --- | --- | --- | --- | --- | --- |
|  |  |  |  |  |  |  |  | Lower | Upper |
| Tigers | Density | 0.01 | 0.01 | 4.26 | 1 | 0.04 | 1.01 | 1.00 | 1.02 |
|  | Noise | 0.04 | 0.03 | 2.61 | 1 | 0.11 | 0.96 | 0.91 | 1.01 |
|  | Constant | 1.30 | 1.58 | 0.68 | 1 | 0.41 | 3.66 | - | - |
| R<sup>2</sup> = 0.01 (Cox & Snell), 0.01 (Nagelkerke); Model $\chi^2$ (2) = 4.17, p < 0.05; Significant at p < 0.05 | | | | | | | | | |
| Leopards | Density | 0.03 | 0.02 | 4.70 | 1 | 0.03 | 1.03 | 1.00 | 1.06 |
|  | Noise | 0.02 | 0.03 | 0.30 | 1 | 0.585 | 0.99 | 0.93 | 1.04 |
|  | Constant | -0.29 | 1.61 | 0.03 | 1 | 0.86 | 0.75 | - | - |
| R<sup>2</sup> = 0.01 (Cox & Snell), 0.02 (Nagelkerke); Model $\chi^2$ (2) = 8.14, p < 0.05; Significant at p < 0.05 | | | | | | | | | |

## 4. Discussion

In response to any change in the environment, alteration of behavior repertoire reflects the first line of defense of an animal. High proportions of inactive behaviors found in the study align with various activity budgeting studies (Biolatti et al., 2016; Mallapur & Chellam, 2002; Pitsko, 2003; Sajjad, Farooq, Anwar, Khurshid, & Bukhari, 2011; Yu et al., 2009). Lack of enrichment elements and hiding refuges in captive conditions may cause excessive inactivity (Mallapur & Chellam, 2002). In this study, all subjects performed the stereotypical behaviors for 7% to 38% of the time in varying forms like pacing, skip-pacing, and tail or toe sucking (Figure 1 and Table 2). Stereotypic pacing is usually accompanied by consistent behavior of marking territory (Boorer, 1972). The high proportion of stereotypy amongst captive felids has been demonstrated in multiple studies (Bashaw et al., 2003; Biolatti et al., 2016; Clubb & Mason, 2007; De Rouck, Kitchener, Law, & Nelissen, 2005; Mallapur, Qureshi, & Chellam, 2002; Mohapatra, Panda, & Acharya, 2014; Sajjad et al., 2011). It has been suggested that the stereotypical level beyond 10% of the total activity is generally unacceptable for any captive animals (Broom, 1983). According to some study, an animal’s welfare status is considered unacceptable if more than 5% of the studied population performs stereotypic behaviors (Mason, 1991; Wielebnowski, 2003). In predatory animals, stereotypies are generally locomotory in nature, which may result from their motivation to forage, range, seek mate, patrol territory, explore, and escape aversive situations (Clubb & Vickery, 2006). The significant pacing levels along enclosure edges were also reported by other studies involving various felid species (Lyons et al., 1997; Mallapur et al., 2002; Sajjad et al., 2011). Intensified pacing and restlessness coincided with feeding and when the food truck was audible or visible to the animals. Bouts of stereotypy also overlapped with the presence of animal keepers around the housing exhibits. Numerous studies on big cats made identical observations (Mohapatra, Mishra, Parida, & Mishra, 2010; Mohapatra et al., 2014; Palita, 1997). The high stereotypies displayed by the two species could be induced by the predictable feeding regime and simplified food provisioning technique. Modification of food, such as hiding it or an unpredictable schedule, can enhance the targeted animal welfare (Shepherdson, Carlstead, Mellen, & Seidensticker, 1993; Watters, Miller, & Sullivan, 2011). Display of stereotypies implies suboptimal welfare in captive conditions as it conveys a warning sign of potential suffering.

This study suggested the increased display of stereotypic behavior in males compare to female conspecifics. Few other studies have reported similar influence of sex on stereotypy in case of captive felids (Dybowska et al., 2008; Vaz et al., 2017). Due to the male big cat’s larger territory size in the wild, the male individuals may experience more spatial stress in enclosed spaces. Tigers bred in captivity exhibited more stereotypy. Similar findings were also reported in another study (Vaz et al., 2017). Adult tigers showed more stereotypic behaviors than young counterparts, as recorded in other studies (Breton & Barrot, 2014; Vaz et al., 2017). This is supported by the motion that stereotypies develop when felids become old enough to disperse from their natal home range and further intensify with age (Mohapatra et al., 2014; Smith, 1993). As animal ages and body enlarges, it experiences spatial constraints in captive conditions, causing behavioral repertoire alterations.

Visitor presence, the noise produced, visitor proximity and behavior, are known to influence captive specie’s behavior repertoire (Hosey & Druck, 1987). Human activities such as shouting, teasing, banging barriers, and throwing stones at the animals may cause psychological and physical harm to the victim animal (Venugopal & Sha, 1993). Visitor effect could induce stress in zoo animals, which may ultimately contribute to the appearance of pathologies and failure of captive breeding programs (Carder & Semple, 2008; Chamove, Hosey, & Schaetzel, 1988; Hosey & Druck, 1987). The study revealed the negative impact of visitor crowd size on the behavior repertoire of tigers and leopards. A large audience crowd has shown to influence stereotypy of captive felid species in various studies (Quadros, Goulart, Passos, Vecci, & Young, 2014; Sellinger & Ha, 2005; Vidal et al., 2016). Leopards housed in enclosures with larger viewing area performed high levels of stereotypic behaviors, thus supporting the effect of visitation on big cats in captivity. Bouts of stereotypy due to visitors presence suggest the animal’s motivation to express flight behavior, but enable to perform the desired behavior (Dembiec, Snider, & Zanella, 2004).

One of the primary purposes of zoological parks is to impart knowledge and the idea of conservation amongst the public. Zoos need to attract visitors and communicate a strong message of conservation to achieve this. Visitor attraction hypothesis suggests that active animals engage visitors more efficiently, while inactive and stereotypic behaviors performed by animals are perceived as boredom and stress by the visitors (Hosey, 2000). Therefore, it is crucial to alleviate the sub-optimal captive conditions that promote stereotypic behaviors amongst big cats. The provision of enrichment techniques is a mean to reduce levels of stereotypy and inactivity in captive felines (Mallapur et al., 2002; Powell, 1995; Skibiel, Trevino, & Naugher, 2007). Provision of cardboard box and toys, hiding refuges, elevated platforms, and olfactory enrichment are few recommended enrichment techniques to ensure optimal welfare (Bashaw et al., 2003; Damasceno et al., 2017; Jenny & Schmid, 2002; Markowitz & LaForse, 1987; McPhee, 2002; Mellen & Shepherdson, 1997; Mohapatra et al., 2010). Such techniques aid to encourage feeding, exploration, and interaction by eliciting species-specific behaviors. Moreover, the installation of appropriate visual barriers between caged animals and visitors is also an efficient measure to reduce the prevalence of stereotypic behaviors (Blaney & Wells, 2004).

## 5. Conclusion and Recommendations

This study suggests that the stereotypic behaviors were prevalent amongst tigers and leopards at National Zoological Park, New Delhi. The levels of stereotypy differed for the biological and captive factors of the big cats. Male, adult, and previously bred individuals exhibited the lengthened pacing periods compared to female and young individuals. Stereotypic behaviors performed by captive tigers and leopards were significantly impacted due to visitation. Large audience size led to an increase in the proportion of time spent in performing stereotypy. Thus, we recommend the installation of visual barriers to minimize the viewing area. Although enclosures at NZP follow the norms laid by the Central Zoo Authority, providing enrichment may possibly reduce the stereotypical behaviors and enhance the captive specie’s welfare. Enrichment elements such as hidden spots and refuges, which mimic the wild, may promote the animals to exhibit more exploratory behaviors.

## Ethics approval

The study involves no animal capture and handling and thus does not require any animal ethics committee permission. However, the necessary permission was obtained from concerned authorities to conduct the study. The subjects were monitored from a distant place without disturbing their natural behavior.

## Acknowledgments

The authorities of the Amity Institute of Forestry and Wildlife, Amity University, and National Zoological Park, New Delhi, are acknowledged for their administrative support and permission to carry out the research work. We wish to thank the zoo staff of National Zoological Park for their assistance during the field study. The study is partially supported by the Bio-bridge Initiative (2019-20) research grant of the Secretariat of the Convention on Biological Diversity and Ministry of Environment of the Republic of Korea.

## Author contribution

AG and PP conceived and designed experiments. RS, UH, and HL improvised the experiment design and assisted in data analysis. AG, SV, and MS collected the data. AG analysed data and wrote the manuscript. All authors read and approved the final version of manuscript.

